# Dopaminergic responses to identity prediction errors depend differently on the orbitofrontal cortex and hippocampus

**DOI:** 10.1101/2024.12.11.628003

**Authors:** Yuji K. Takahashi, Zhewei Zhang, Thorsten Kahnt, Geoffrey Schoenbaum

## Abstract

Adaptive behavior depends on the ability to predict specific events, particularly those related to rewards. Armed with such associative information, we can infer the current value of predicted rewards based on changing circumstances and desires. To support this ability, neural systems must represent both the value and identity of predicted rewards, and these representations must be updated when they change. Here we tested whether prediction error signaling of dopamine neurons depends on two areas known to represent the specifics of rewarding events, the HC and OFC. We monitored the spiking activity of dopamine neurons in rat VTA during changes in the number or flavor of expected rewards designed to induce errors in the prediction of reward value or reward identity, respectively. In control animals, dopamine neurons registered both error types, transiently increasing firing to additional drops of reward or changes in reward flavor. These canonical firing signatures of value and identity prediction errors were significantly disrupted in rats with ipsilateral neurotoxic lesions of either HC or OFC. Specifically, HC lesions caused a failure to register either type of prediction error, whereas OFC lesions caused persistent signaling of identity prediction errors and much more subtle effects on signaling of value errors. These results demonstrate that HC and OFC contribute distinct types of information to the computation of prediction errors signaled by dopaminergic neurons.

## Introduction

Learning to predict outcomes is critical to adaptive behavior. The content of such learning includes information about an outcome’s value at the time of learning and information about its defining characteristics or features, such as its sensory properties. While much is known about the neural systems mediating the former, far less is understood about the machinery supporting the latter. Yet this information is critical, since only by knowing these features can we flexibly update our predictions to reflect the current outcome value or utility at the time of choice (Behrens et al., 2018). Such inference cannot be done if we only store a pointer to the static value at the time of learning.

To solve this problem, we need neural systems that predict these features combined with teaching signals to update their predictions when features change. Such sensory, identity, or feature-specific prediction errors could be similar to so-called reward or value prediction errors, only operating across dimensions other than value. Accordingly, we have reported that the midbrain dopamine neurons, which famously signal value prediction errors (Dabney et al., 2020; Eshel et al., 2015; Glimcher, 2011; Mirenowicz and Schultz, 1994; Schultz et al., 1997; Waelti et al., 2001), also respond to value neutral changes in the identity of an expected reward in both rats and humans (Howard and Kahnt, 2018; Suarez et al., 2019; Takahashi et al., 2017). Consistent with the proposal that such activity may act as teaching signals to support learning about reward features, these firing changes are not general but instead encode the conjunction between the error and the identity of the unexpected reward (Howard et al., 2024; Stalnaker et al., 2019), and dopamine transients are both necessary and sufficient for learning to predict specific rewards (Chang et al., 2017; Keiflin et al., 2019b; Steinberg et al., 2013).

If dopamine neurons signal errors in predicting features of reward beyond value (Gershman et al., 2024; Kahnt and Schoenbaum, 2025), as captured in a variety of new models (Gardner et al., 2018; Lee et al., 2022; Millidge et al., 2023; Takahashi et al., 2023), then key upstream regions critical for the associated behaviors should be involved in this process. In line with this prediction, recent work has shown that updates to outcome representations in the human OFC are correlated with the strength of identity prediction errors in the midbrain (Howard and Kahnt, 2018), and network-targeted TMS designed to disrupt these representations in OFC causes persistent error signaling in midbrain in response to reward identity shifts (Qingfang et al., 2024).

Here we built on this work, recording midbrain dopamine neurons in rats performing a task in which we independently manipulated reward value or reward identity to induce prediction errors. Neural correlates of these errors in controls were compared to those in rats with ipsilateral hippocampal or orbitofrontal lesions. The results demonstrate that signaling of identity prediction errors at the time of reward by midbrain dopamine neurons depends on unique contributions from both areas.

## Results

We recorded single-unit activity in the VTA from rats with ipsilateral sham (n = 5) or neurotoxic lesions of HC (n = 9) or OFC (n = 9) (see figure 2 for surgical coordinates). In the HCx group, lesions targeted the entire HC, resulting in visible loss of neurons in 54% (49-60%) of this region across subjects. In the OFCx group, lesions targeted the medial and lateral part of OFC, resulting in visible loss of neurons in 50% (30 – 63%) of this region across subjects (Fig. 1).

**Figure 1:**
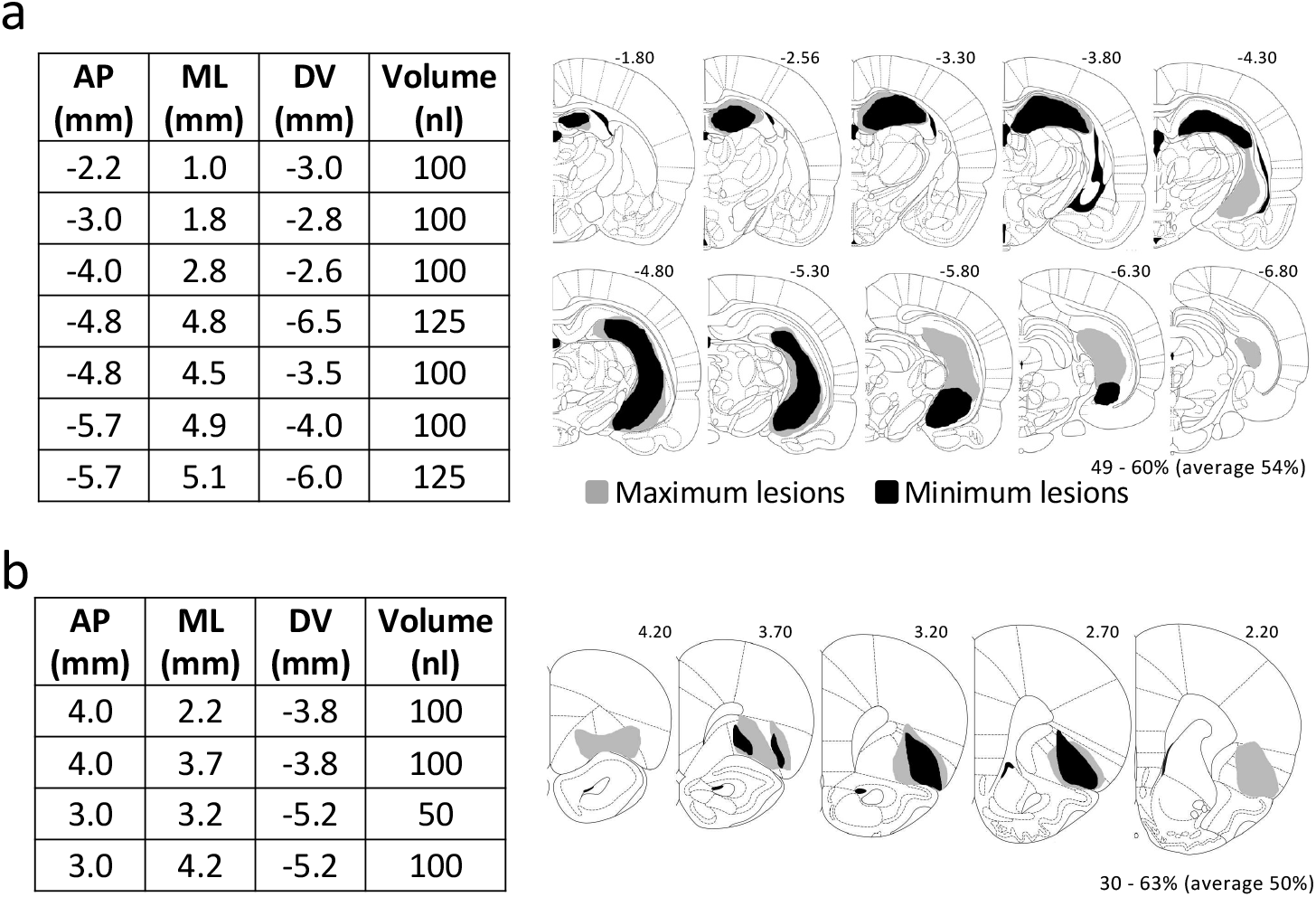
Surgical coordinates and extent of ipsilateral hippocampal and orbitofrontal lesions. **(a and b)** Table gives volumes and coordinates (AP and ML relative to bregma and DV relative to brain surface) of injections. Brain sections illustrate the extent of the maximum (gray) and minimum (black) lesion at each level in HCx (a) and OFCx (b) in the lesioned rats.

**Figure 2:**
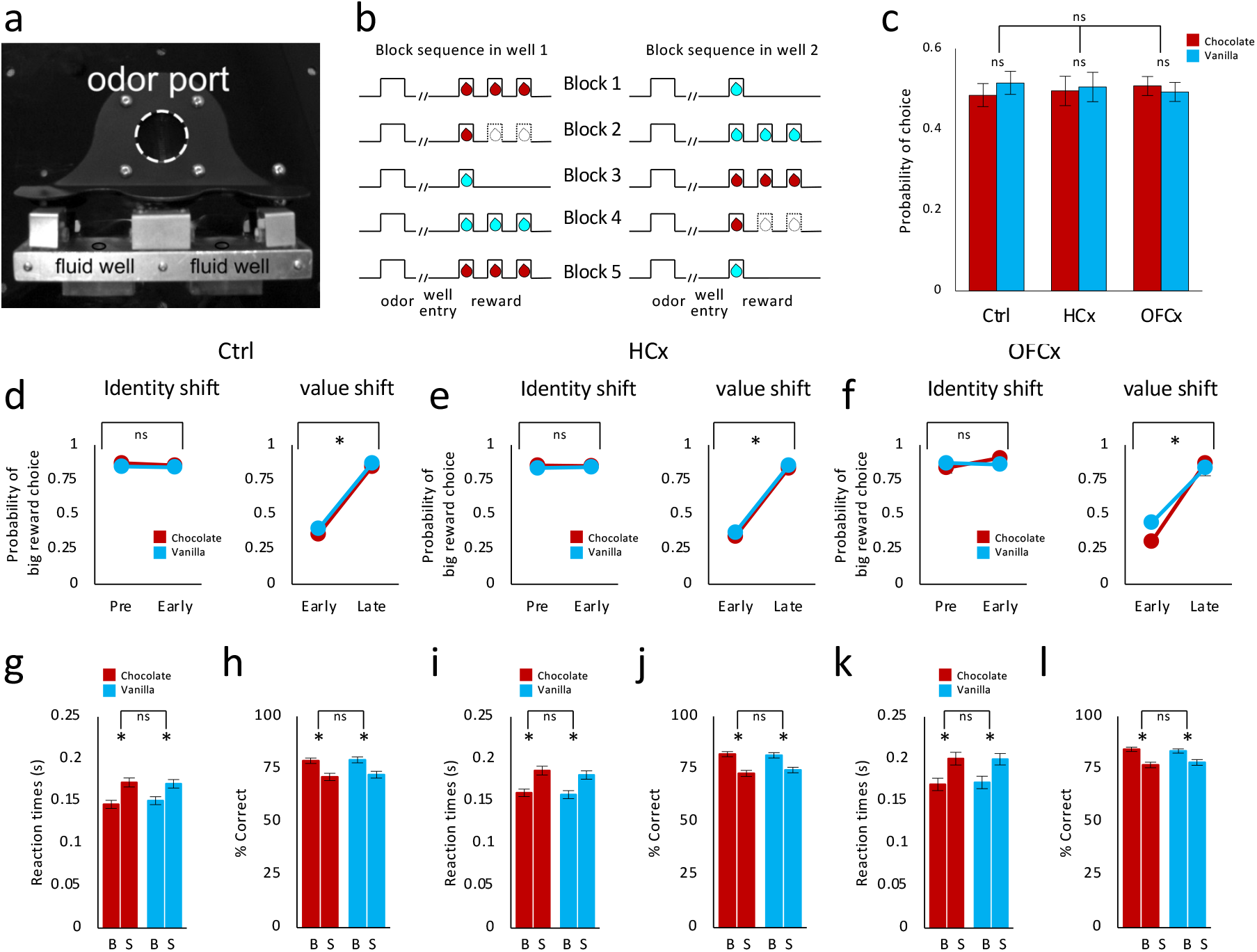
Task design and behavior. **(a)** Picture of apparatus used in the task, showing the odor port (∼2.5 cm diameter) and two fluid wells. **(b)** Deflections indicate the time course of stimuli (odors and rewards) presented to the rat on each trial. Dashed lines show when a reward was omitted, and solid lines show when reward was delivered. At the start of each recording session, one well was randomly designated to deliver the big reward, which consisted of three drops of flavored milk (chocolate or vanilla). One drop of the other flavored milk was delivered in the other well (block 1). In the second and fourth blocks, the number off drops delivered in the two wells was switched without changing the flavors (value shift). In the third and fifth blocks, the flavors delivered in the two wells were switched without changing the number of drops (identity shift).In all three groups, chocolate- and vanilla-flavored milk was equally preferred in two-flavor choice test (F’s <0.28, p’s > 0.62) and there were no significant main effects or interactions with group (F’s < 0.19, p’s > 0.78). **(d – f)** Effect of value and identity switches on choice behavior in Control (d), HCx (e) and OFCx (f) groups. Across identity shifts there were no effects of block and no interactions with either group or identity (F’s < 2.02, p’s > 0.16), whereas across value shifts, there were significant main effects of block overall and within each group (F’s > 344.0, p’s < 0.000) and no interactions with either identity or group (F’s < 3.1, p’s > 0.08). **(g-l)** Reaction time and % correct in response to high and low valued reward of each identity on forced choice trials in control (g), HCx (i) and OFCx (k) groups. All groups showed faster reaction times and higher accuracy when the high valued reward was at stake (F’s > 31.0, p’s < 0.0000), and there were no interactions with either identity or group (F’s < 1.6, p’s > 0.19).

Neurons were recorded in rats performing an odor-guided identity shift task (Fig. 2a-b), which was previously used to characterize signaling of prediction errors across shifts in reward value versus reward flavor (Takahashi et al., 2017). One each trial, rats sampled one of three different odor cues at a central port and then responded at one of two adjacent fluid wells (Fig. 2a) to receive a big (three drop) or a small (one drop) amount of one of two equally preferred flavored milk solutions (chocolate or vanilla, Fig. 2c). One odor signaled the availability of reward only in the left well (forced left), a second odor signaled reward only in the right well (forced right), and a third odor signaled that reward was available at both (free choice). To induce errors in reward prediction, we manipulated either the value or the flavor of the rewards produced by each well across blocks of trials (Fig. 2b).

Analysis of behavior during recording confirmed that rats in all groups tracked the rewards available at each well across blocks, as expected based on previous work showing that unilateral lesions to OFC or HC do not impair animals’ ability to respond to changes in reward on this simple task (Takahashi et al., 2011; Zhang et al., 2023). Across both flavors, rats in all three groups consistently chose the larger reward more often on free-choice trials (Figs 2d-f) and responded with shorter reaction times and more accurately on forced-choice trials when the larger reward was at stake (Figs. 2g-l). ANOVAs revealed main effects of reward number (F’s > 31.0, p’s < 0.001) and no interactions involving group or identity (F’s < 3.1, p’s > 0.08).

Dopamine neurons recorded during these sessions were identified by means of a waveform analysis, previously shown to identify TH+ neurons (Takahashi et al., 2017). Using this approach, we categorized as putative dopamine neurons 80 of 408 VTA neurons in controls, 110 of 513 VTA neurons in HCx, and 90 of 412 VTA neurons in OFCx rats (Fig. 3), proportions which did not differ between groups (Chi-square = 0.71, p > 0.05). Of these, 46, 63 and 47 increased firing to reward in controls, HCx and OFCx rats, respectively (t-test, p < 0.05, compared with a 400 ms baseline taken during the inter-trial interval before trial onset).

**Figure 3:**
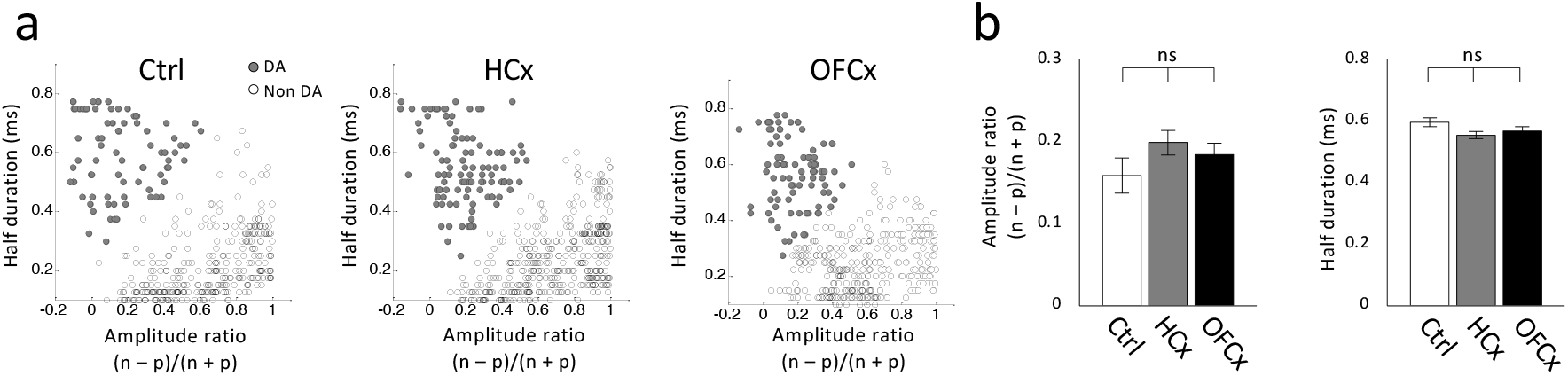
Identification and waveform features of putative dopamine neurons. **(a)**Result of cluster analysis based on the half-time of the spike duration and the ratio comparing the amplitude of the first positive and negative waveform segments ((n – p) / (n + p)). Data in left show VTA neurons (n = 408) from the control group, plotted as putative dopamine neurons (gray circles, n = 80), and neurons that classified with other cluster (open circles, n = 328). Data in left show VTA neurons (n = 513) from the HCx group, plotted as putative dopamine neurons (gray circles, n = 110), and neurons that classified with other cluster (open circles, n = 403). Data in right show VTA neurons (n = 412) from the OFCx group, plotted as putative dopamine neurons (gray circles, n = 90), and neurons that classified with other cluster (open circles, n = 322). **(b)** bar-graphs indicate average amplitude ratio (left) and half duration (right) of putative dopamine neurons in control (open), HCx (gray) and OFCx (black) groups. ANOVA revealed no significant main effect of group in amplitude ratio (F_2,277_ = 1.44, p > 0.24) and half duration (F_2,277_ = 2.59, p > 0.07).

### Ipsilateral hippocampal and orbitofrontal lesions disrupt error signaling in response to changes in reward number

As expected, the prediction error signaling induced by changes in reward number was evident in the reward-responsive dopamine neurons in controls. These neurons changed firing in response to shifts in reward number at the start of blocks 2 and 4, increasing to unexpected delivery and decreasing to unexpected omission of the 2^nd^ drop of reward (Fig. 4a). These changes in firing were maximal immediately after a block switch and diminished with learning in later trials (Fig. 4b; statistics in captions). To further quantify these changes in firing, we computed difference scores for each neuron, comparing the average firing on early vs late trials after the number shift at the time of delivery or omission of 2^nd^ drop. Distributions of these scores were shifted above zero when an unexpected reward was delivered (Fig. 4g, left) and below zero when an expected reward was omitted (Fig. 4g, right).

**Figure 4:**
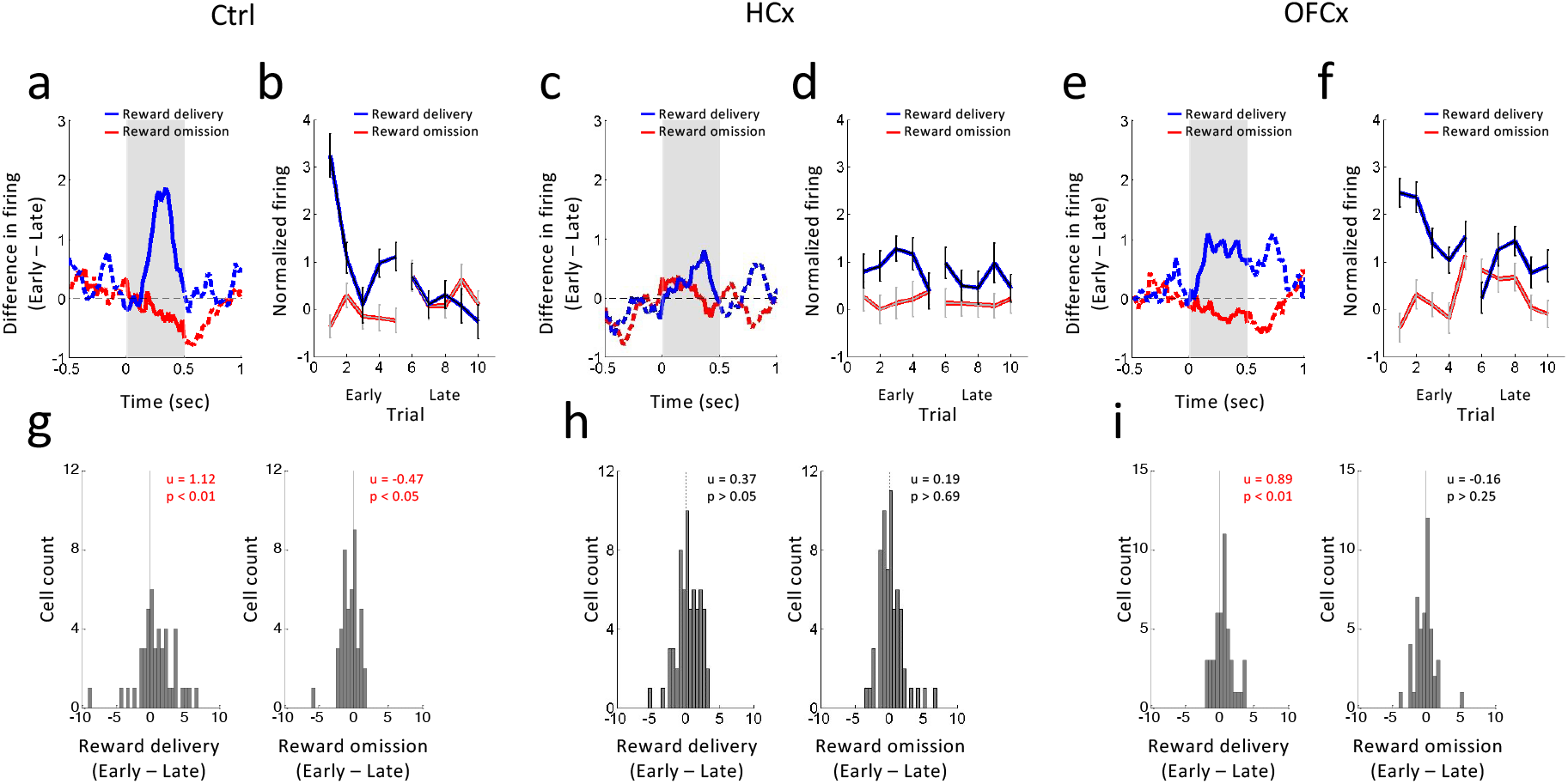
Changes in reward-evoked activity of reward-responsive dopamine neurons to changes in reward value. **(a, c and e)** Population responses show the difference in firing between early and late trials in response to unexpected reward delivery (blue) and unexpected omission (red) in control (a), HCx (c), and OFCx (e) groups. Gray shadings indicate the time when the reward was manipulated. **(b, d and f)** Average firing in response to reward delivery (blue) and omission (red) in the first 5 and last 5 trials in control (b), HCx (d) and OFCx (f) groups. Error bars, S.E.M. ANOVA (Reward x Early/Late x Trial) revealed significant main effects of Reward in all groups (Control, F_1,87_ = 7.47, p<0.01; HCx, F_1,117_ = 8.28, p<0.01; OFCx, F_1,89_ = 21.2, p<0.01), and significant interactions between Reward x Early/Late in Control (F_1,87_ = 13.1, p<0.01) and OFCx groups (F_1,89_ = 12.5, p<0.01). Planned contrasts revealed a significant main effect of Early/Late for reward delivery in Controls (F_1,87_ = 9.78, p<0.01) and OFCx (F_1,89_ = 10.6, p<0.01), but not in HCx (F_1,117_ = 1.18, p>0.05) and a main effect of Early/Late for reward omission in control (Control, F_1,87_ =4.47, p<0.05), but not in HCx (F_1,117_ = 0.08, p=0.78) or OFCx (F_1,89_ = 1.28, p=0.26). **(g – i)** Distributions of difference scores comparing firing to unexpected reward delivery (left) and omission (right) in the early and late trials in control (g), HCx (h) and OFCx (i) groups. The numbers in each panela indicate results of Wilcoxon singed-rank test (p) and the average difference score (u).

As previously reported (Takahashi et al., 2011; Zhang et al., 2023), these firing changes were disrupted by ipsilateral lesions of HC or OFC. This was most marked in HCx rats where reward-responsive dopamine neurons failed to change firing when a reward was unexpectedly delivered or omitted (Fig. 4c; statistics in caption), resulting in difference scores comparing firing early vs late that were not different from zero (Fig. 4h). By contrast, ipsilateral OFC lesions had a more subtle effect on signaling of reward prediction errors. While neurons in the OFCx rats increased firing when an unexpected reward was delivered and then declined (Fig. 4e-f; statistics in caption), the contrast was less impressive than in controls (F1,293 = 5.77, p = 0.01). Additionally, these neurons failed to exhibit a significant decrease in firing when an expected reward was omitted (Fig. 4e-f; statistics in captoin). Consistent with this impression, difference scores comparing activity early vs late in these neurons shifted above zero to unexpected reward (Fig. 4i, left) but showed no change from zero when reward was omitted (Fig. 4i, right).

### Ipsilateral hippocampal and orbitofrontal lesions disrupt error signaling in response to changes in reward identity

We previously reported that reward-responsive dopamine neurons increased firing in response to value-neutral changes in reward identity (Takahashi et al., 2017). To test whether such signals were affected by ipsilateral lesions of HC or OFC, we compared firing of dopamine neurons before and after shifts in reward identity in blocks 3 and 5. As expected, reward-responsive dopamine neurons in controls increased firing to each drop of both the big and small reward when the identity was changed unexpectedly (Fig. 5a). These increases in firing were generally maximal immediately after identity shifts, and then diminished on later trials (Fig. 5b; statistics in caption). We again quantified these changes in firing by computing difference scores for each neuron, in this case comparing firing to reward at the start of the new block versus the end of the previous block. These scores shifted above zero in response to the 1^st^ (Fig. 5g, left) and 2^nd^ drops (Fig. 5g, middle) of the big reward and to the single drop of the small reward (Fig. 5g, right).

**Figure 5:**
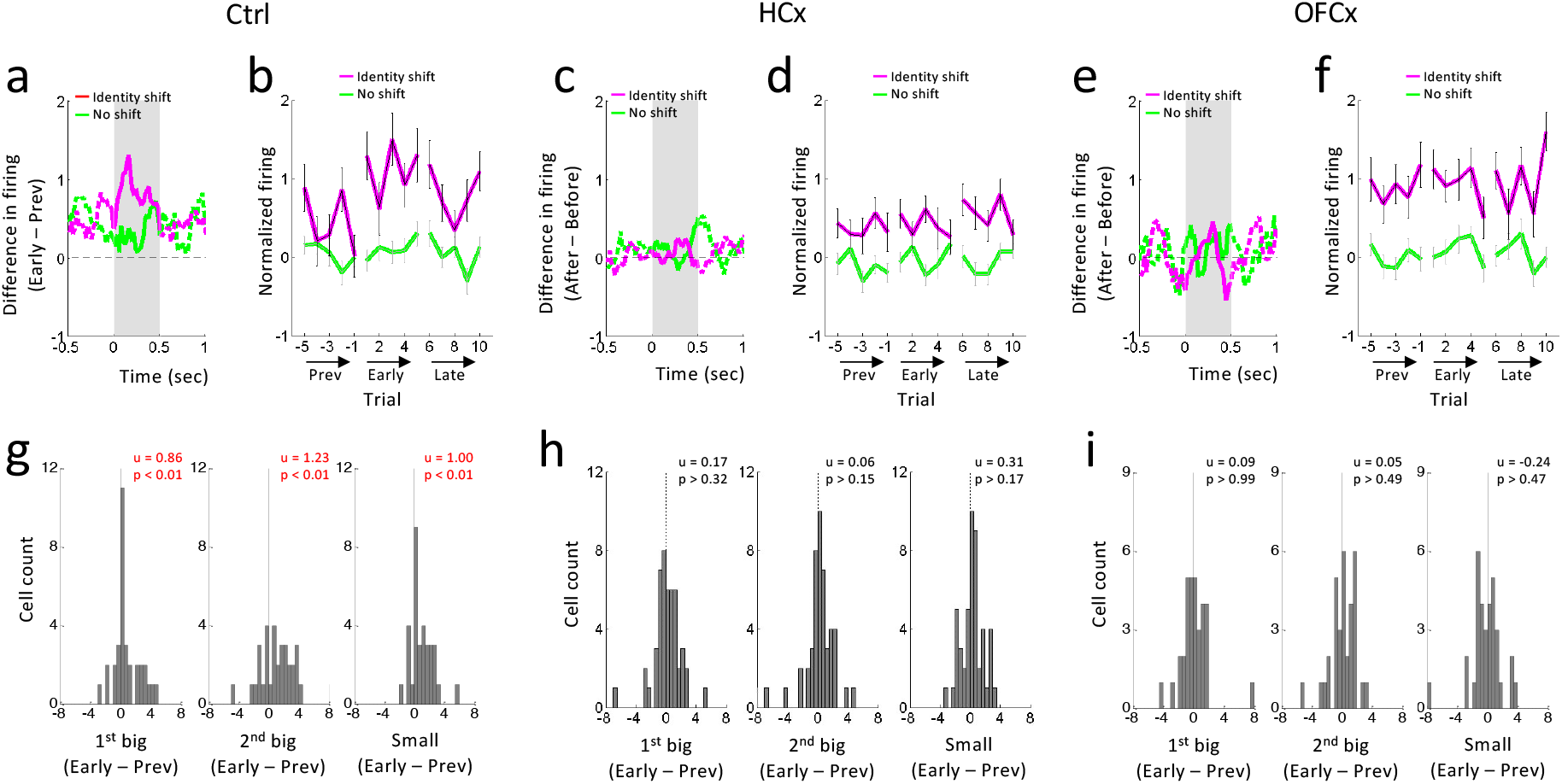
Changes in reward-evoked activity of reward-responsive dopamine neurons to changes in reward identity. **(a, c and e)** Difference in population activity between last 5 trials before and first 5 trials after a shift of reward identity for average of 1^st^ , 2^nd^ big reward and small reward (purple) and a comparison period where there was no shift taken 0.5 and 1.0s after small reward (green) in control (a), HCx (c) and OFCx (e) groups. Gray shadings indicate the time when the reward was manipulated. **(b, d and f)** Changes in average firing before and after reward identity shift in control (b), HCx (d) and OFCx (f) groups. Purple lines indicate average firing at the time of the 1^st^ and 2^nd^ drops of the big reward and the small reward. Green lines indicate average firing in comparison period after small reward. Error bars, SEM. ANOVA (Shift/No shift x Prev/Early/Late x Trial) revealed a significant main effect of Shift/No shift in all groups (Control, F_1,45_ = 15.3, p < 0.01; HCx, F_1,64_ = 16.1, p < 0.01; OFCx, F_1,42_ = 18.9, p < 0.01), and a significant interaction between Shift/No shift x Prev/Early/Late in Control (F_2,90_ = 4.74, p < 0.05), but not in HCx and OFCx (F’s < 0.76, p’s > 0.47). Planned contrasts revealed a significant main effect of Prev/Early/Late in firing after shifts in Control (F_2,90_ = 6.40, p < 0.01), but not in HCx (F_2,128_ = 0.71 p > 0.49) and OFCx (F_2,84_ = 0.08, p > 0.92). There were no significant main effects or interactions in firing in the comparison period in any group (F’s < 72, p’s >0.49). **(g – i)** Distributions of difference scores comparing firing to first (left), second (middle) drops of big reward and small reward (right) in the last 5 versus first 5 trials before and after an identity shift in control (g), HCx (h) and OFCx (i) groups. The numbers in each panela indicate results of Wilcoxon singed-rank test (p) and the average difference score (u).

By contrast, dopamine neurons recorded in both groups of lesioned rats failed to increase firing to the new reward flavor after identity shifts, instead maintaining relatively consistent levels of firing (Fig 5c-f; statistics in caption). As a result, difference scores for neurons recorded in both groups of rats were distributed around zero to the 1^st^ (Figs. 5h and 5i, left) and 2^nd^ drops (Figs. 5h and 5i, middle) of the big reward and to the single drop of the small reward (Figs. 5h and 5i, right). Interestingly, however, dopamine neurons in HCx rats maintained low baseline-normalized activity across the identity shift (Fig. 5d), while dopamine neurons in OFCx rats maintained high levels of activity (Fig. 5f). This apparent difference was confirmed by ANOVAs comparing firing early versus later across groups. These analyses revealed significant interactions between controls and both OFCx (F1,151 = 5.21, p = 0.02) and HCx rats (F1,151 =7.07, p = 0.008) but not between OFCx and HCx rats (F1,151 = 0.02, p = 0.88), where only a main effect of group was evident (F1,151 = 4.44, p = 0.03). Direct comparisons revealed that the activity in HCx rats was similar to that late in blocks in controls (p’s > 0.11), whereas activity in the OFCx rats was similar to that on early trials in controls (p’s > 0.45). In other words, while shifts in reward identity briefly caused transient increases in dopamine neuron activity in controls, those same changes drove persistent increases in dopamine neuron activity in rats with OFC lesions while having relatively little impact on dopamine activity in rats with HC lesions.

## Discussion

Here we monitored the spiking activity of dopamine neurons in rat VTA during changes in the number or flavor of expected rewards designed to induce errors in the prediction of reward value or reward identity, respectively. In controls, dopamine neurons registered both error types, transiently increasing firing to additional drops of reward (and suppressing firing upon reward omission) as well as to changes in reward flavor. In rats with ipsilateral neurotoxic lesions of either HC or OFC, these firing patterns were significantly disrupted, but in different ways. HC lesions caused a failure to register either type of prediction error, whereas OFC lesions caused persistent signaling of identity prediction errors and much more subtle effects on signaling of value errors.

In considering these data, it is important to emphasize that many aspects replicate prior work, in both rats and humans. This begins with the demonstration of effects of OFC and HC lesions on the signaling of value prediction errors. Lesioning or inactivation of OFC has been shown to attenuate dopamine neuron firing to unexpected reward and to eliminate the suppression of firing to reward omission (Jo et al., 2013; Jo and Mizumori, 2016; Takahashi et al., 2011), whereas HC lesions prevent changes in firing to either (Zhang et al., 2023).

The current data also replicate prior work in rats showing that dopamine neurons that normally respond to changes in value also register value-neutral changes in reward flavor (Takahashi et al., 2017), a result that has also been extended to humans (Howard et al., 2024; Howard and Kahnt, 2018). While initially controversial, this finding is now incorporated into novel models using temporal difference frameworks that attempt to explain dopaminergic error signaling beyond value (Gardner et al., 2018; Lee et al., 2022; Millidge et al., 2023; Takahashi et al., 2023), and it can account for otherwise hard-to-explain evidence that transient changes in the activity of midbrain dopamine neurons is both necessary and sufficient to support learning about specific reward features (Chang et al., 2017; Gardner et al., 2018; Keiflin et al., 2019a; Steinberg et al., 2013). Finally, in a parallel task in humans, we have reported that network-targeted TMS designed to disrupt OFC activity caused extended signaling of identity prediction errors after shifts, remarkably similar to the persistent error signaling caused by complete OFC lesions here (Qingfang et al., 2024). The consistency of these aspects of the data across studies, labs, and species increases confidence in their reliability and also the reliability of the new findings.

In this regard, the demonstration that prediction error signals reflecting changes in reward identity depend on both HC and OFC is particularly notable, since although identity error signals have been incorporated into new temporal difference algorithms (Chang et al., 2017; Gardner et al., 2018; Keiflin et al., 2019a; Steinberg et al., 2013), these stop short of recognizing a role for dopaminergic errors in true model-based processing (Langdon et al., 2017). Given the prominent position of OFC and HC in most circuits devoted to model-based learning and behavior (Whittington et al., 2019; Wikenheiser and Schoenbaum, 2016; Wilson et al., 2014; Zeithamova et al., 2012), this position seems increasingly untenable, unless one holds that even these areas are engaged in something less than model-based processing (Momennejad et al., 2017; Stachenfeld et al., 2017).

The differential effects of OFC and HC lesions on error responses are also notable, since they show these two areas make distinct contributions, at least to VTA dopamine neuron error signaling. To recap, while ipsilateral lesions of both structures prevented changes in firing in response to identity switches, the HC lesions resulted in low activity similar to that *after* learning in controls, whereas OFC lesions resulted in high activity similar to that *before* learning in controls. Thus, without HC, dopamine neurons are blind to changes in reward identity, whereas without OFC, they are constantly surprised.

The apparent blindness of dopamine neurons in the HC-lesioned hemisphere to violations of identity expectations is similar to the effect observed to violations of value expectations, where low signals were also observed. This similarity suggests a single common function in both situations, explained by modeling work in our prior work as reflecting a loss of strong priors, such that changes in rewards update priors rather than resulting in prediction errors and new learning (Zhang et al., 2023). This explanation holds here for identity errors, and further suggests that HC is not necessary for making the predictions themselves but rather provides temporal or mnemonic support for them, at least at the time of reward. By contrast, the persistence of identity prediction errors in dopamine neurons in the OFC-lesioned hemisphere, also observed in humans after TMS affecting OFC (Qingfang et al., 2024), is more consistent with an area that is a critical source of these predictions, since failed or attenuated updating (previously shown to correlate with the strength of error signaling (Howard and Kahnt, 2018)) would result in ongoing errors.

Further, OFC seems to be preferentially important for identity predictions and not value predictions, since OFC lesions did not cause persistent value-based prediction errors, either here or in our prior study. This is of course contrary to the strong version of the hypothesis that the OFC is the seat of all value predictions (Levy and Glimcher, 2011; Padoa-Schioppa and Assad, 2006; Padoa-Schioppa and Conen, 2017), and instead suggests that some other area – perhaps striatum (Daw et al., 2005; Joel et al., 2002; Yin et al., 2004) – can generate much of this information for downstream consumption without the top-down input from OFC. This conclusion is more consilient with data showing that OFC is not generally necessary for value-guided behavior (Chudasama et al., 2007; Gardner et al., 2017; Noonan et al., 2010; Schoenbaum et al., 2002), except during initial learning, when training has been relatively limited, or when new information must be integrated with established knowledge (Gardner et al., 2019; Gardner et al., 2020; Gore et al., 2023; Kuwabara et al., 2020). The idea that OFC is more critical to outcome information than information about value is also in accord with numerous causal studies implicating OFC in outcome-specific behavior (Izquierdo and Murray, 2007; Lichtenberg et al., 2017; McDannald et al., 2014; McDannald et al., 2005; Ostlund and Balleine, 2007; Pickens et al., 2005; Rudebeck et al., 2013; Tegelbeckers et al., 2023).

## Materials availability

This study did not generate unique reagents.

## Data and code availability

All data reported in this manuscript will be archived and made publicly available as of the date of publication as per NIH regulations and the lead contact will provide data upon request and assist other researchers with analysis and interpretation.

## Acknowledgments

This work was supported by the Intramural Research Program at the National Institute on Drug Abuse (Z1A-DA000587). The opinions expressed in this article are the authors’ own and do not reflect the view of the NIH/DHHS. The authors have no conflicts of interest to report.

## Author Contributions

YKT, TK, and GS designed the experiment, and YKT conducted the experiment (surgery, behavioral training, unit recording, and post-mortem tissue processing). YKT analyzed the resultant data, with input from GS, ZZ, and TK, and YKT, ZZ, TK, and GS collaborated to interpreted the data and write the manuscript.

## STAR Methods

### Lead contact

Requests for further information and resources should be directed to and will be fulfilled by Dr. Geoffrey Schoenbaum.

## Experimental model and subject details

### Subjects

Twenty-three male Long-Evans rats (Charles River Labs, Wilmington, MA) were used in this study. Rats were tested at the NIDA-IRP in accordance with NIH guidelines. Fourteen of them (5 rats in control and 9 rats in HCx) were also part of a previous study in which dopamine neurons were recorded in a different task (Zhang et al., 2023).

## Method details

### Stereotaxic Surgery

All surgical procedures adhered to guidelines for aseptic technique. For recording, a drivable bundle of eight 25-um diameter formvar insulated nichrome wires (A-M systems, Carlsborg, WA) was chronically implanted dorsal to VTA in the left or right hemisphere at 5.3 mm posterior to bregma, 0.7 mm laterally, and 7.5 mm ventral to the brain surface at an angle of 5° toward the midline from vertical. In addition, rats received neurotoxic lesion of ipsilateral hippocampus (n = 9) or orbitofrontal cortex (n = 5) by infusion of 20 mg/ml NMDA (see Fig 1 for the surgical coordinates). Controls (n = 5) received sham lesions in which burr holes were drilled and the pipette tip lowered into the brain but no solution delivered. Cephalexin (15 mg/kg p.o.) was administered twice daily for two weeks post-operatively.

### Histology

All rats were perfused with phosphate-buffered saline (PBS) followed by 4% paraformaldehyde (Santa Cruz Biotechnology Inc., CA), after which the brains were removed, cut in 40 µm sections, and stained with thionin to visualize lesions and electrode tracks.

### Odor-guided choice task

Recording was conducted in aluminum chambers approximately 18” on each side with sloping walls narrowing to an area of 12” x 12” at the bottom. A central odor port was located above two fluid wells (Fig. 2a). Two lights were located above the panel. The odor port was connected to an air flow dilution olfactometer to allow the rapid delivery of olfactory cues. Odors were chosen from compounds obtained from International Flavors and Fragrances (New York, NY). Trials were signaled by illumination of the panel lights inside the box. When these lights were on, a nosepoke into the odor port resulted in delivery of the odor cue to a small hemicylinder located behind this opening. One of three different odors was delivered to the port on each trial. At odor offset, the rat had 3 seconds to make a response at one of the two fluid wells. One odor instructed the rat to go to the left well to get reward, a second odor instructed the rat to go to the right well to get reward, and a third odor indicated that the rat could obtain reward at either well. Odors were presented in a pseudorandom sequence such that the free-choice odor, indicating reward was available at either well, was presented on 7/20 trials, while the forced choice odors were presented on 6 trials each. In addition, the same odor could be presented on no more than 3 consecutive trials. Once the rats were shaped to perform this basic task, we introduced blocks in which we independently manipulated the size or the identity of the reward (Fig. 2b). At the start of each recording session, one well was randomly designated to deliver the big reward in which three boli of chocolate or vanilla milk were delivered, and the other to deliver the small reward in which one bolus of the other flavored milk was delivered (Block 1, Fig. 2b). The bolus size was ∼0.05 ml and the time between boli was 500 ms. In the second block of trials, the reward value was switched by changing the number of drops without changing flavors. (Block 2, Fig. 2b). In the third block of trials, the reward identity was switched by changing the flavor of the reward without changing the number of drops (Block 3, Fig. 2b). The value and identity switches were repeated one more time in Block 4 and Block 5, respectively (Fig. 2b). Block 1 was 30-50 trials long and all subsequent blocks were 60-100 trials long. Additionally block switches required rats to have chosen the large reward on at least 6 of the last 10 free-choice trials. Sessions in which rats did not complete all blocks were not included in the analyses.

### Flavor preference testing

To test for flavor preferences in the task, we also ran several sessions for each rat, consisting entirely of free choice trials, in which one well produced 2 drops of chocolate milk and the other well produced 2 drops of vanilla milk, switching location three times across 4 blocks of trials. We assessed free choice performance during these blocks in order to identify any preference between the two flavored rewards without confounding value shifts.

## Quantification and statistical analysis

### Single-unit recording

Wires were screened for activity daily; if no activity was detected, the rat was removed from the box, and the electrode assembly was advanced 40 or 80 µm. Otherwise, active wires were selected to be recorded, a session was conducted, and the electrode was advanced at the end of the session. Neural activity was recorded using Plexon Multichannel Acquisition Processor systems (Dallas, TX). Signals from the electrode wires were amplified 20X by an op-amp headstage (Plexon Inc, HST/8o50-G20-GR), located on the electrode array. Immediately outside the training chamber, the signals were passed through a differential pre-amplifier (Plexon Inc, PBX2/16sp-r-G50/16fp-G50), where the single unit signals were amplified 50X and filtered at 150-9000 Hz. The single unit signals were then sent to the Multichannel Acquisition Processor box, where they were further filtered at 250-8000 Hz, digitized at 40 kHz and amplified at 1-32X. Waveforms (>2.5:1 signal-to-noise) were extracted from active channels and recorded to disk by an associated workstation.

### Data analysis

Units were sorted using Offline Sorter software from Plexon Inc (Dallas, TX). Sorted files were then processed and analyzed in Neuroexplorer and Matlab (Natick, MA). Dopamine neurons were identified via a waveform analysis. Briefly a cluster analysis was performed based on the half time of the spike duration and the ratio comparing the amplitude of the first positive and negative waveform segments. The center and variance of each cluster was computed without data from the neuron of interest, and then that neuron was assigned to a cluster if it was within 3 s.d. of the cluster’s center. Neurons that met this criterion for more than one cluster were not classified. This process was repeated for each neuron. The putative dopamine neurons that showed increase in firing to reward (100-500 ms after delivery) compared to baseline (400ms before reward) were further classified as reward-responsive (t-test, p< 0.05).

